# Structure of the Human Secretory Immunoglobulin M Core

**DOI:** 10.1101/2020.09.10.291138

**Authors:** Nikit Kumar, Christopher P. Arthur, Claudio Ciferri, Marissa L. Matsumoto

## Abstract

Immunoglobulins (Ig) A and M are the only human antibodies that form oligomers and undergo transcytosis to mucosal secretions via the polymeric Ig receptor (pIgR). When complexed with the J-chain (JC) and the secretory component (SC) of pIgR, secretory IgA and IgM (sIgA and sIgM) play critical roles in host-pathogen defense. Recently, we determined the structure of sIgA-Fc which elucidated the mechanism of polymeric IgA assembly and revealed an extensive binding interface between IgA-Fc, JC, and SC. Despite low sequence identity shared with IgA-Fc, IgM-Fc also undergoes JC-mediated assembly and binds pIgR. Here, we report the structure of sIgM-Fc and carryout a systematic comparison to sIgA-Fc. Our structural analysis reveals a remarkably conserved mechanism of JC-templated oligomerization and SC recognition of both IgM and IgA through highly a conserved network of interactions. These studies reveal the structurally conserved features of sIgM and sIgA required for function in mucosal immunity.

## Introduction

The human immune system is composed of five major classes of immunoglobulins (Ig). IgG, IgD, and IgE exist only as monomers, whereas IgA and IgM have the ability to form polymers due to an 18 amino acid tailpiece extension on their heavy chains (Sørensen et al., 1996). IgA dimers, tetramers, and pentamers can form only in the presence of the J-chain (JC), a 15 kDa, cysteine-rich polypeptide required for its polymeric assembly (Johansen et al., 2000). In contrast, polymeric IgM can form both pentamers and hexamers, depending on the presence or absence of the JC, respectively (Heyman and Shulman, 2016; Kownatzki and Drescher, 1973).

Hexameric IgM is best known for its role in complement activation through the classical pathway via C1q (Sharp et al., 2019), whereas pentameric IgM plays an important role in host pathogen defense at the mucosa (Johansen et al., 1999). Like polymeric IgA, JC-containing, pentameric IgM binds to the polymeric Ig receptor (pIgR) and can be transcytosed across the epithelium to carry out its protective function in mucosal tissues (Johansen et al., 2000). Upon transcytosis the pIgR is proteolytically cleaved releasing the ectodomain of the receptor known as the secretory component (SC), which remains associated with IgA and IgM (Johansen et al., 2000). The released Ig-SC complex is referred to as secretory IgA or IgM (sIgA or sIgM). While sIgA is the predominant immunoglobulin found in mucosal secretions, sIgM plays an important role, particularly highlighted by the fact that selective IgA deficiency does not typically result in recurrent infections due to the ability of sIgM to provide protection (Catanzaro et al., 2019).

Despite extensive study, the structures of sIgA and sIgM have remained elusive for many years. Recently we determined the first atomic-resolution structures of dimeric, tetrameric, and pentameric assemblies of sIgA-Fc (Kumar et al., 2020). Our structures revealed the novel fold of the JC, which acts as a template for IgA-Fc oligomerization mediated by the tailpieces. The interaction of the JC with the IgA-Fcs imparts asymmetry on the IgA oligomer allowing one-to-one binding with the SC. The SC binds across the IgA oligomer making extensive interactions with both the JC and Fcs. Furthermore, the structures revealed that the SC undergoes a large conformational rearrangement from a closed, apo state (Stadtmueller et al., 2016), to an open, extended conformation upon polymeric IgA binding. Our structures are highly consistent with two additional reported structures of murine (https://doi.org/10.1101/2020.02.16.951780) and human sIgA (Wang et al., 2020).

To better understand whether the mechanism of polymerization and SC recognition is conserved across both secreted Ig isotypes, we determined the structure of the human sIgM-Fc core by cryo-electron microscopy (cryo-EM) and carried out a detailed comparative analysis with the structure of sIgA-Fc. Although slight differences in the flexibility of the JC are observed along with minor differences in binding by the SC, the architecture of sIgA and sIgM is highly homologous and consistent with the recently reported structure of sIgM by Li and colleagues (Li et al., 2020). Our structures reveal a mechanism of JC-mediated oligomerization that is remarkably conserved in both sIgA and sIgM. Preservation of the molecular interactions required for SC recognition between IgA and IgM suggests highly similar pIgR binding and transcytosis mechanisms for both secreted Igs. These studies reveal the highly conserved nature of secreted polymeric Ig assembly, structure, and function.

## Results

### Cryo-EM Structure of the Secretory Immunoglobulin M Core

Prior analysis of recombinant full-length pentameric IgM by negative stain EM revealed significant flexibility of the Fab arms (Hiramoto et al., 2018). In order to obtain a rigid assembly for structural determination, full-length IgM was truncated to IgM-Fc containing only the heavy chain constant domains (Cμ2, Cμ3, and Cμ4) and the tailpiece. IgM-Fc was co-expressed with the JC to obtain a pentameric assembly. To further rigidify the pentamers, a complex was formed with the secretory component (SC) of the pIgR and single particle cryo-EM was used to obtain a 3D reconstruction (Fig. 1 – figure supplement 1 and Table 1). The sample displayed significant preferential orientation, which was overcome by collecting additional data at a tilt angle of 40 degrees. Data from zero and 40 degree tilt were processed to yield a 3.25 Å reconstruction [Fourier shell correlation (FSC) = 0.143] of the complex that included the Cμ3, Cμ4, and tailpiece segments of the IgM-Fc, the JC, and the SC. Despite Cμ2 being included in the construct, flexibility of this domain precluded structure determination. Regions of the map containing the highest local resolution were at the complex core and included the tailpiece segments as well as the interface between the SC domain 1 (D1), IgM Cμ4, and the JC (Fig. 1 – figure supplements 2 and 3). Homology models of the human IgM Cμ3, Cμ4, and tailpieces were built based on the NMR structure of murine Cμ3 (PDB code 4BA8), the crystal structure of murine Cμ4 (PDB code 4JVW), and the tailpiece segments from pentameric sIgA-Fc (PDB code 6UEA), respectively. These models along with the JC and individual domains of the SC from pentameric sIgA-Fc (PDB code 6UEA) were docked, rigid-body fit into the map, and refined.

**Table 1:**
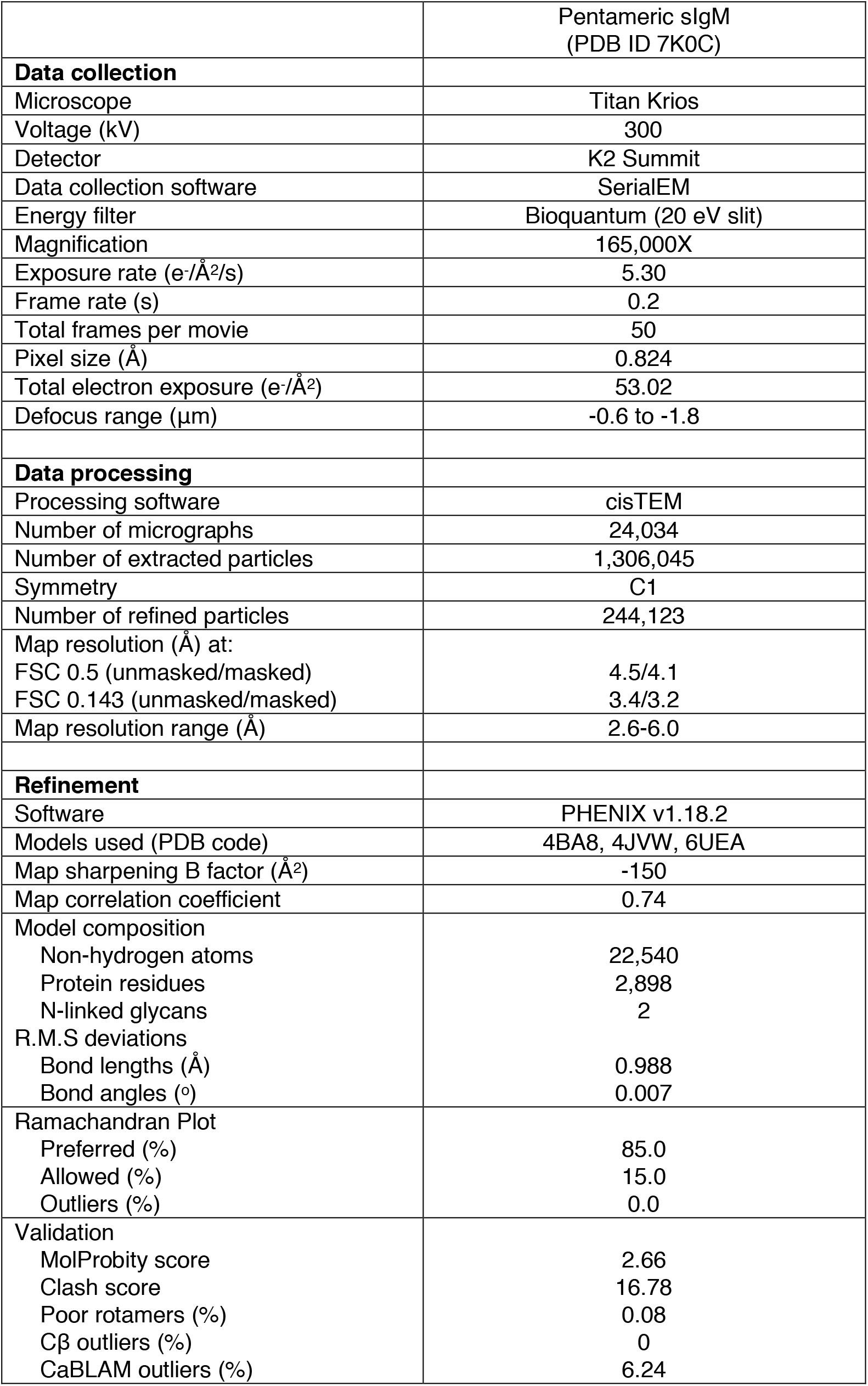
CryoEM data collection, refinement, and validation statistics.

**Figure 1:**
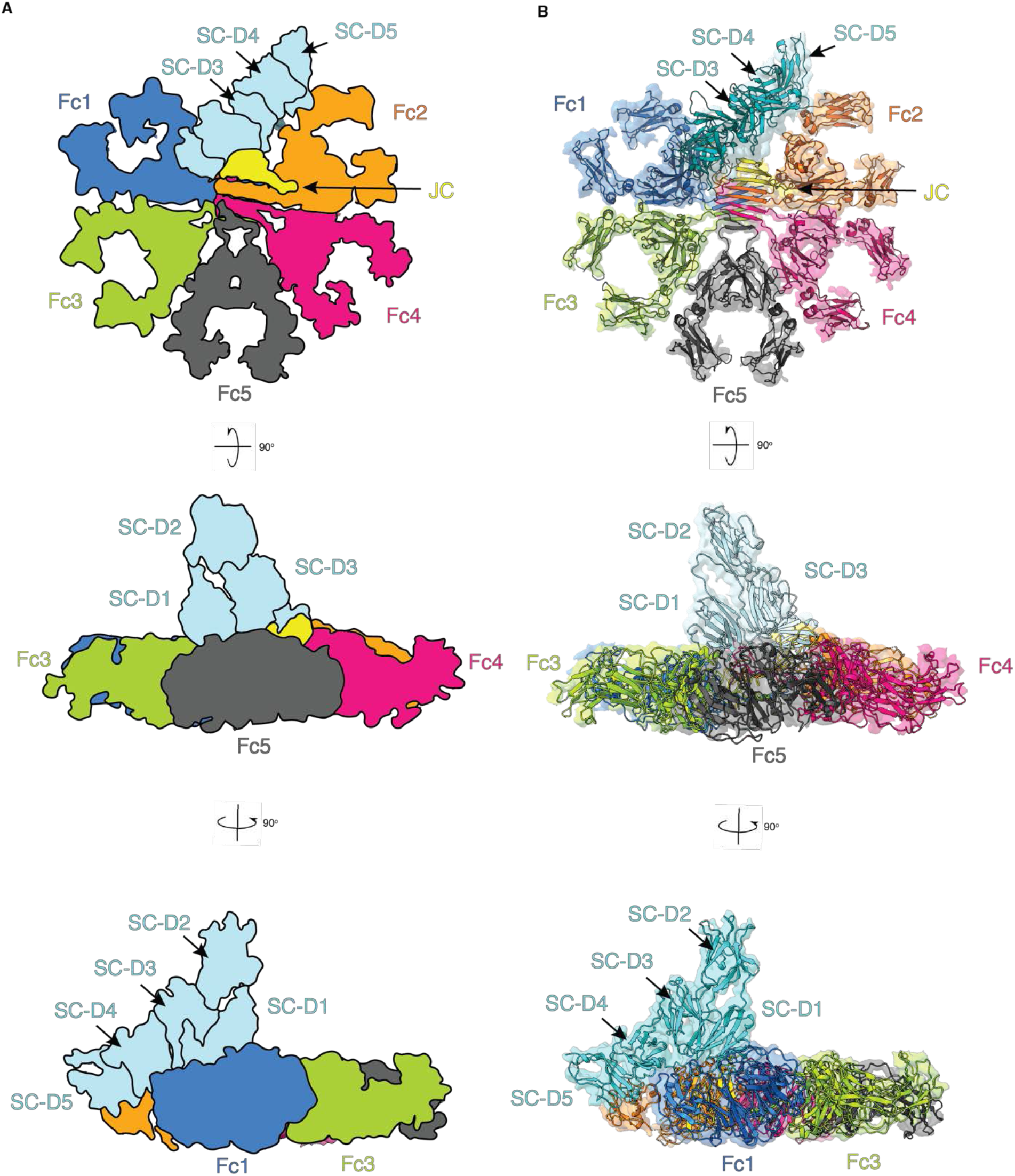
Cryo-EM structure of the pentameric sIgM complex. (A) Top, front, and side schematics of the sIgM complex. (B) Transparent cryo-EM maps overlaid with the model are shown for the complex as in (A).

Overall, sIgM forms an asymmetric pentamer, with the J-chain occupying the site where a sixth IgM-Fc would be expected in a symmetric, hexameric assembly (Fig. 1). This observation is in line with previously reported negative stain analysis of IgM-Fc complexed with JC and the apoptosis inhibitor of macrophage molecule (AIM) (Hiramoto et al., 2018). The five Fcs lie in-plane, stabilized by inter-Fc disulfide bonds between C414 in the Cμ3 domain of adjacent Fcs, as suggested previously (Davis et al., 1989; Sørensen et al., 1999). Additionally, the Fcs are arranged radially around a central β-sandwich composed of the JC and the tailpieces of the five Fcs. The SC binds across the top face of the pentamer in a bent, but extended conformation, contacting Fc1, Fc2 and the JC. The overall architecture of sIgM-Fc is strikingly reminiscent of pentameric sIgA-Fc ((Kumar et al., 2020), PDB code 6UEA).

### Structure of the J-chain

With an overall resolution of 3.25 Å and the highest local resolution at the core of the IgM-Fc/JC/SC interaction, the JC model from our previous sIgA structure could be easily docked into the map. A major difference between the sIgA and sIgM structures is that in sIgM, we did not observe density for residues 71-98 of the 137-residue JC (Fig. 2A). This is consistent with the structure reported by Li and colleagues (Li et al., 2020). In IgA, as previously characterized, the JC interacts with the Cα3 domains of IgA through its three β-hairpin regions, with β-hairpins 1 and 2 interacting on the top surface of Fc2, while β-hairpin 3 interacts with the bottom surface of Fc1 (Fig. 2B). While β-hairpins 1 and 3 are similarly observed to interact with the Cμ4 domains of IgM to the top and bottom surfaces of Fc2 and Fc1, respectively, residues 71-98 corresponding to β-hairpin 2 were unstructured (Fig. 2A). JC β-hairpin 1 interacts with IgM via the packing of JC_I22_ and JC_V34_ with IgM Fc2 and anchored by a salt-bridge interaction between JC_E31_ with IgM Fc2_R461_ (Fig. 2C). In pentameric IgA, JC_I22_ and JC_V34_ similarly pack against IgA Fc2, while the tip of the β-hairpin is anchored by a salt-bridge interaction between JC_D32_ and IgA Fc2_R450_ (Fig. 2C). Interestingly, while IgM_R461_ is not conserved in IgA (Fig. 2 – figure supplement 1A), the salt-bridge interaction is maintained by a neighboring residue, IgA_R450_, located near this interface. Also, instead of forming the salt-bridge with JC_E31_, IgA instead uses the adjacent residue JC_D32_. JC β-hairpin 3 primarily packs residues JC_V114, L116, Y118, V125_ against hydrophobic IgM residues Fc1_L359, F485, V537, V547_ in a groove formed between Cμ3 and Cμ4 of IgM Fc1 (Fig. 2D). This interaction is similar to the interaction of JC β-hairpin 3 with IgA where JC_V114, L116, Y118, V125_ pack against homologous IgA residues Fc1_L258, M433, F443_ in a hydrophobic groove between domains Cα2 and Cα3 of IgA Fc1 (Fig. 2D). With the exception of β-hairpin 2, which is unstructured in sIgM, the structure of the JC is very similar between pentameric sIgM and sIgA complexes, with an overall RMSD of 1.30 Å (Fig. 2E).

**Figure 2:**
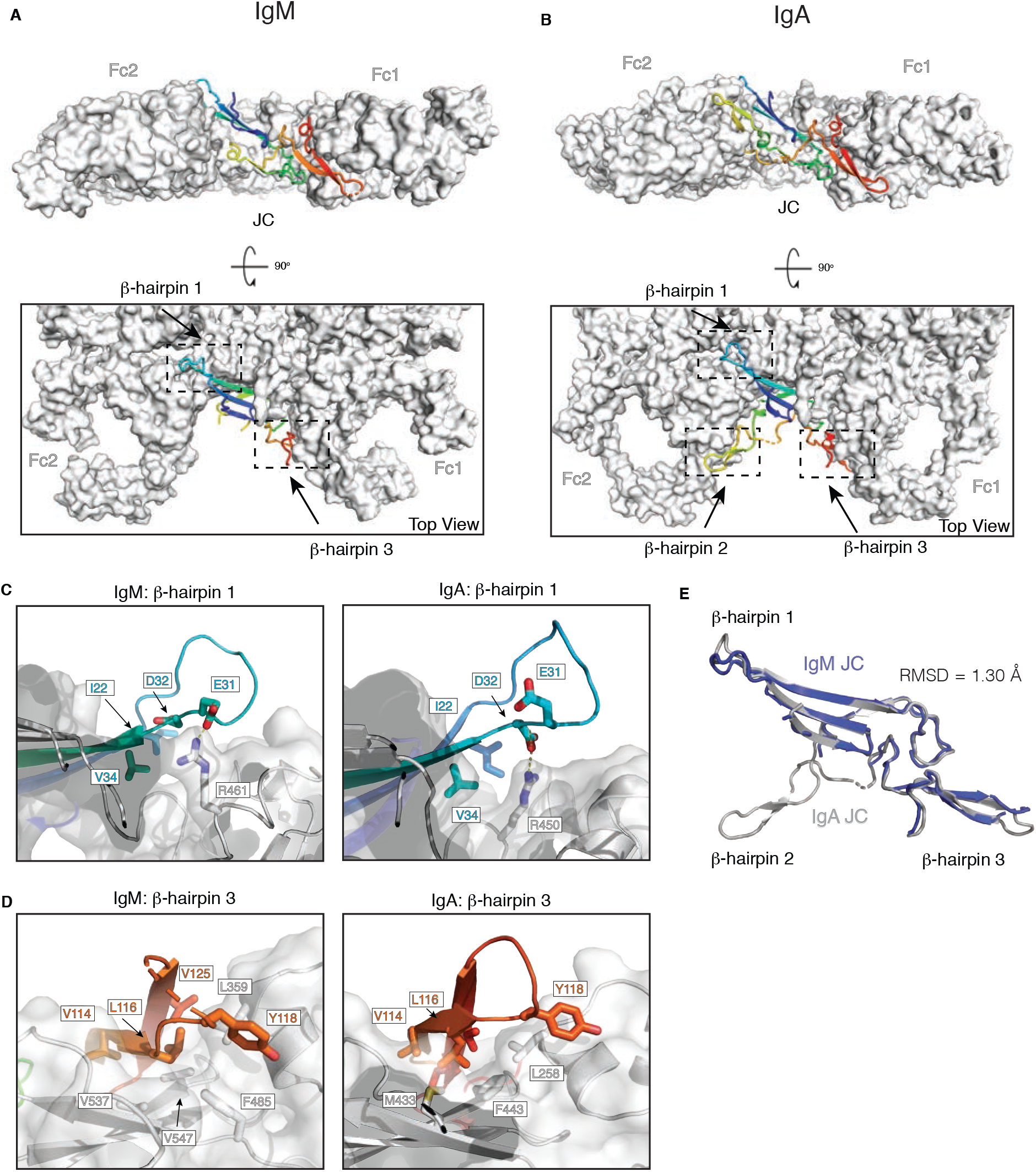
Structural similarities of the JC assembled with pentameric IgM and IgA. (A) Back and top views of the JC in pentameric sIgM colored in rainbow from the N-terminus in blue to the C-terminus in red. The SC has been omitted for clarity. (B) Identical views and coloring of the J chain as in (A) for pentameric sIgA. (C) Magnified views of JC β-hairpin 1 from sIgM (left) and sIgA (right) along with contacts to the Fcs. (D) Magnified views of J chain β-hairpin 3 from IgM (left) and IgA (right) along with contacts to the Fcs. (E) Alignment of the JC from sIgM in blue to the JC from sIgA in gray shows an RMSD of 1.30 Å as measured after by structural superposition across 104 residues. (B,C,D) Modified from Kumar et al., 2020 with permission from AAAS.

### Mechanism of IgM Polymerization

In addition to clasping Fc1 and Fc2 of IgM, the JC also templates a central β-sandwich structure that is extended by the tailpieces of each IgM Fc. Tailpiece residues L561 to S569 from Fc1 and Fc3 contribute a pair of parallel β-strands each that extend the bottom face of the central β-sandwich, while Fc2 and Fc4 contribute a pair of parallel β-strands each that extend the top face (Fig. 3A). Meanwhile, Fc5 contributes one β-strand to both the top and bottom of the central β-sandwich and effectively caps this assembly, resulting in a β-sandwich structure containing parallel β-strands on each surface with an anti-parallel arrangement of the top and bottom sheets. This assembly is identical to the central β-sandwich assembly of the IgA pentamer (Fig. 3B). Despite minor differences in individual amino acid composition of the tailpieces (Fig. 2 – figure supplement 1A), the arrangement results in a striking array of repeated residues across the β-sandwich, with primarily hydrophobic residues packed in the center (including IgM Fc_Y562, V564, L566, M568_ as compared to IgA Fc_I458, V460, V462, M464_), while hydrophilic residues remain solvent exposed (including IgM Fc_N563, S565_ as compared to IgA Fc_H457, N459, S461_) (Fig. 3C-D).

**Figure 3:**
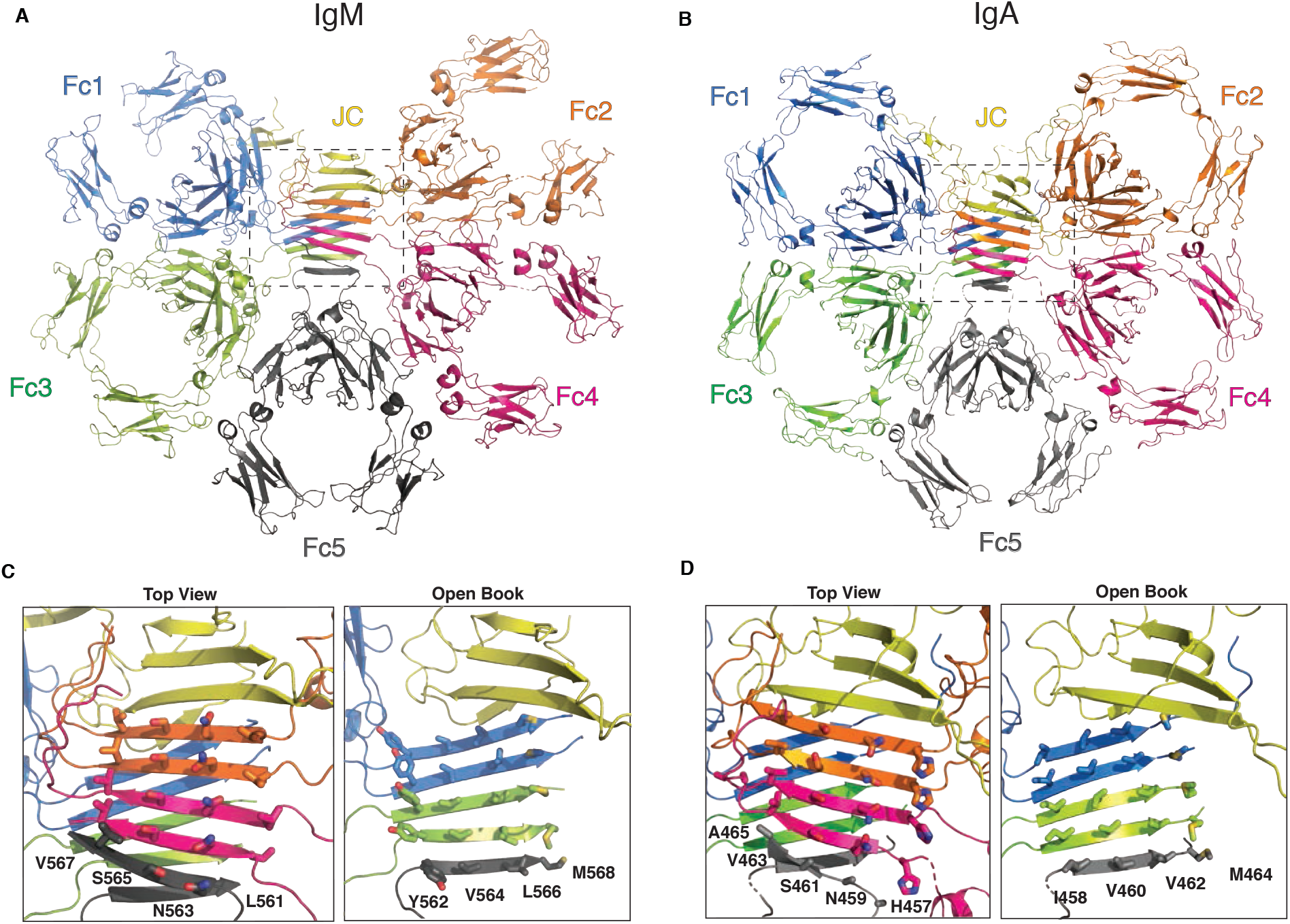
Comparison of tailpiece segments in the oligomerization of pentameric IgM and IgA. Top views of (A) pentameric IgM and (B) pentameric IgA represented without the SC. (C) Magnification of IgM boxed region in (A) with solvent facing, repeating residues on the top surface of the tailpiece segments of the Fcs (left) and open book view with inward facing, repeating residues labelled and shown as sticks. (D) Similar representation as in (C) for IgA with repeating residues labelled and shown as sticks on top surface (left) and inner surface (right). (B,D) Modified from Kumar et al., 2020 with permission from AAAS.

### Recognition of IgM by the Secretory Component

The transport of IgM across epithelial cells requires the recognition by the pIgR. The SC of the pIgR is composed of five Ig-like domains (D1 – D5) (Stadtmueller et al., 2016), and previous biochemical investigations established the involvement of the three CDR-like loops of D1 as necessary for interaction with JC-containing, polymeric IgM and IgA (Coyne et al., 1994; Klimovich, 2011). In the absence of IgM or IgA, the SC adopts a triangular, closed conformation where the CDR residues of D1 are partially or completely buried in an interface between the D1-D4-D5 domains (Stadtmueller et al., 2016). When bound to pentameric IgM, SC D2-D5 are placed head-to-tail in a linear arrangement, while the linker between D1-D2 is turned nearly 180 ° to position D1 for interaction with IgM Fc1 (Fig. 4A). SC-D1 is responsible for the bulk of the interactions with the IgM Fcs and the JC through its three CDR-like loops (Fig. 4B), while SC-D5 packs against IgM Fc2. Interestingly, while a disulfide bond is clearly formed between SC_C468_ and IgA Fc2_C311_ in sIgA, weak density between SC_C468_ and the homologous residue in IgM, Fc2_C414_ was observed at high contour levels of the map, suggesting partial oxidation of the disulfide in our sIgM structure. A covalent interaction between the SC and IgM/JC assemblies has been previously reported (Longet et al., 2012).

**Figure 4:**
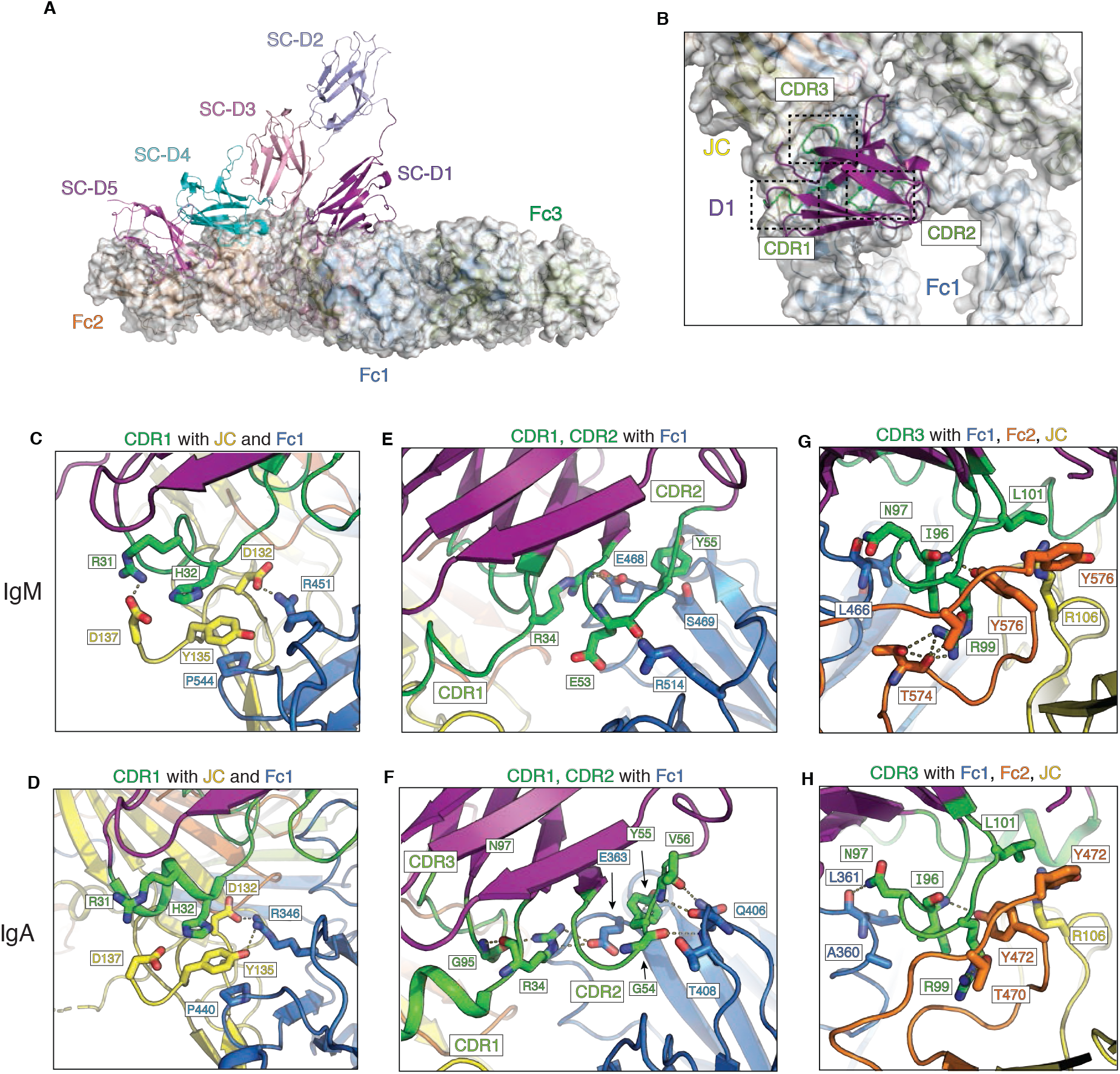
Recognition of pentameric IgM and IgA by SC. (A) sIgM pentamer shown as colored model overlaid with white surface representation of Fcs and JC. Individual domains D1 – D5 of the SC are colored differentially. (B) Top-down view of SC-D1 and its interactions with IgM-Fc/JC through its three CDR regions (boxed and colored green). D1-CDR1 interactions with JC and Fc1 for IgM (C) and IgA (D). Interactions of D1-CDR1 and -CDR2 with Fc1 of IgM (E) and IgA (F). Interactions of D1-CDR3 with Fc1, Fc2, and JC of IgM (G) and IgA (H). Side chains are shown as sticks and polar interactions shown as dashes. (D,F,H) Modified from Kumar et al., 2020 with permission from AAAS.

D1 of the SC recognizes pentameric IgM and IgA through a network of conserved interactions involving all three of its CDR-like loops. In sIgM, CDR1 interacts via a salt-bridge between D1_R31_ and JC_D137_ along with cation-pi stacking interactions between D1_H32_ with JC_Y135_, which in turn packs favorably against IgM Fc1_P544_. Additionally, JC_D132_ forms a salt-bridge with IgM Fc1_R451_ (Fig. 4C). The mechanism of CDR1 recognition is conserved in sIgA with identical JC interactions (with a slightly different rotamer conformation for D1_R31_) and packing against the homologous IgA Fc1_P440_ and salt-bridging with Fc1_R346_ (Fig. 4D). D1_R34_ additionally forms a salt-bridge with IgM Fc1_E468_ (Fig. 4E), which is conserved in sIgA with Fc1_E363_ (Fig. 4F). CDR2 makes backbone interactions between D1_E53_ and IgM Fc1_R514_, along with favorable packing of D1_Y55_ against Fc1_S469_ (Fig. 4E). While these exact CDR2 interactions are not conserved in sIgA, similar compensatory interactions are formed. Backbone interactions occur between D1_G54_ and D1_V56_ with IgA Fc1_T408_ and Fc1_Q406_, respectively, along with favorable stacking of D1_Y55_ against Fc1_E363_ (Fig. 4F), such that the overall conformation of CDR2 is similar in sIgA and sIgM. CDR3 makes a complex network of interactions involving residues from Fc1, Fc2, and the JC in both IgM and IgA complexes. Specifically, the backbone amine of SC_I96_ forms a hydrogen-bond with IgM Fc2_Y576_, SC_N97_ hydrogen-bonds with the backbone carbonyl of IgM Fc1_L466_, while SC_R99_ interacts with IgM Fc2_T574_ and sandwiches Fc2_Y576_ with JC_R106_ (Fig. 4G). This mechanism of interaction with IgA is highly conserved, with analogous interactions between SC_I96_ with IgA Fc2_Y472_, SC_N97_ with Fc1_L361_, and SC_R99_ with IgA Fc2_T470_, Fc2_Y472_, and JC_R106_ (Fig. 4H). Overall, the SC-D1 interaction interface in sIgM and sIgA is highly conserved, despite sharing an overall sequence identity of only 40% between the two Fcs (Fig. 2 – figure supplement 1A).

## Discussion

Secreted polymeric Igs play a critical role in host-pathogen defense processes at mucosal surfaces. In this study, we report the structure of the human sIgM-Fc core, comprising IgM Fc domains Cμ3 and Cμ4, JC, and the SC, and assess the structural similarities with pentameric sIgA. Despite sharing just 40% sequence identity, both immunoglobulins are able to incorporate the JC and transcytose to mucosal surfaces by interacting with the pIgR. A detailed structural comparison between pentameric sIgM and sIgA reveals that conserved key interaction interfaces with the JC and the SC allows for a remarkably similar assembly of these secreted Igs. These include the association of JC β-hairpins-1 and -3 with the Cα2-Cα3 and Cμ3-Cμ4 domains, the Fc tailpiece-mediated extension of a central β-sandwich, and a conserved interface for the interaction with SC-D1. During the preparation of this manuscript, a 3.4 Å-resolution cryo-EM structure of sIgM-Fc was also published (Li et al., 2020). Our independent study reveals a highly consistent structure, further validating these novel findings.

While the structures of sIgA and sIgM are highly similar, we were unable to observe residues corresponding to JC β-hairpin 2 in sIgM, which is also consistent with the observations of Li and colleagues (Li et al., 2020). This was unexpected given the interaction interface between JC β-hairpin 2 and IgA-Fc primarily consists of van der Waals interactions and the hydrophobic nature of the structurally homologous residues are generally conserved in IgM-Fc. Perhaps minor charge differences caused by the replacement of IgA_L441_ to IgM_N545_ at the base of the β-hairpin 2-binding site or insufficient shape complementarity at the IgM-Fc Cμ3-Cμ4 interface are responsible. Despite this difference, the remaining JC structure is very similar to that adopted in pentameric sIgA with an RMSD of 1.30 Å.

The conformation of the SC when bound to IgM-Fc or IgA-Fc is nearly identical, with the majority of interactions mediated by the CDR regions of SC-D1, as predicted from mutagenesis studies (Kaetzel, 2005). Interestingly, while we had previously observed clear density for a single disulfide bond linking SC-D5_C468_ to IgA-Fc_C311_, we observed weak density for a disulfide between D5_C468_ and IgM-Fc_C414_ despite using an identical protocol for complex formation containing reduced glutathione. Previous reports have indicated that the SC is covalently attached to IgM (Longet et al., 2012), while other reports indicate that SC non-covalently interacts with IgM (Hamburger et al., 2004; Stadtmueller et al., 2016). Based on our observations where SC-D5 orients in a similar conformation when complexed with either IgA-Fc or IgM-Fc, along with the close spatial proximity of the respective cysteines, it is likely that there was a mixture of particles in our sIgM-Fc sample that either contain the disulfide or those that are non-covalently interacting.

Overall, our study provides structural insights into the pentameric assembly of IgM-Fc complexed with the JC and bound to the SC. Both polymeric IgM and IgA complexed with JC also interact with Fcα/μR with likely overlapping binding sites as SC-D1 (Hamburger et al., 2004), while only IgM interacts with the FcμR/Faim3/Toso (Liu et al., 2019). Further structural characterization of the interaction with these receptors would allow for a better understanding of the mechanism of receptor specificity for IgM and provide further insights into the role of IgM beyond mucosal immunity.

## Materials and Methods

### Construct Generation

The protein sequence for human IgM-Fc was obtained from UniProt (www.uniprot.org). A sequence encoding residues 226-576 was synthesized and cloned into a mammalian pRK expression vector containing an N-terminal signal sequence, FLAG-tag, and PreScission protease cleavage site. Constructs encoding human JC (residues 23-159) and human SC (residues 19-603 containing a C-terminal hexahistidine tag) was described previously (Kumar et al., 2020; Lombana et al., 2019).

### Protein Expression and Purification

Vectors encoding IgM-Fc and JC were transiently co-transfected into CHO DP12 cells as previously described (Wong et al., 2010) in a 4:1 DNA mass ratio. Media containing secreted IgM-Fc/JC assemblies was purified by anti-FLAG affinity resin (Genentech, South San Francisco, CA), washed with buffer A (50 mM Tris, pH 7.5, 150 mM NaCl, 5 mM EDTA, and 2 mM NaN_3_), eluted with 50 mM sodium citrate, pH 3.5, 150 mM NaCl, and neutralized with 0.2 M arginine, 137 mM succinate, pH 9.0. SC was similarly transfected into CHO DP12 cells and purified as described (Kumar et al., 2020; Wong et al., 2010). Briefly, media containing secreted SC was applied to Ni Sepharose Excel resin (GE Healthcare, Chicago, IL), washed with buffer B (50 mM NaPO4, pH 7.4, 200 mM naCl, and 1 mM NaN_3_), and eluted with buffer B supplemented with 400 mM Imidazole. Fractions containing IgM-Fc/JC complexes or SC were concentrated with a 100- or 30-kDa MWCO Amicon centrifugal filtration device, respectively, prior to loading to a HiLoad Superdex 200 pg 26/60 column (GE Healthcare) equilibrated with buffer C (20 mM HEPES, pH 7.2, 250 mM NaCl). Peak fractions corresponding to pentameric IgM-Fc/JC complex or SC were recovered and concentrated.

### CryoEM Sample Preparation and Data Collection

Purified IgM-Fc/JC assemblies at 1 uM were complexed with 2-fold molar excess of SC in the presence of 0.3 mM reduced Glutathione (Sigma, St. Louis, MO) and incubated overnight at 4°C. IgM-Fc/JC/SC complexes were separated from excess SC via size exclusion chromatography using a Superose 6 increase 3.2/300 (GE Healthcare) in Buffer C described above. Peak fractions corresponding to complexes were collected and diluted to 0.2 mg/mL. Prior to cryo-grid preparation, samples were briefly cross-linked with 0.05% glutaraldehyde (Sigma) for 10 mins at 20 °C. Cryo-grids were prepared as previously described (Kumar et al., 2020). Briefly, Quantifoil R 0.6/1 holey carbon grids were freshly glow discharged for 5 s prior with a Solarus 950 Plasma Cleaner (Gatan, Pleasanton, CA) prior to application of 3 uL protein sample. Grids were blotted with Vitrobot Filter Paper (Electron Microscopy Sciences, Hatfield, PA) for 2.5 s with a blot force of 8 using a Vitrobot Mark IV (Thermo Fisher, Waltham, MA) set to 4°C with 100% humidity, and plunged into liquid nitrogen-cooled liquid ethane. Movie stacks were collected on a Titan Krios (Thermo Fisher) operating at 300 kV equipped with K2 Summit (Gatan) direct electron detector using SerialEM (Mastronarde, 2005). Images were collected at 165,000X calibrated to a pixel size of 0.824 Å/pixel. Movie stacks containing 50 images were recorded every 0.2 s with a total electron exposure of 53.02 electrons/Å^2^.

### Data Processing

18,168 movie stacks at zero-degree tilt were collected, along with 5,866 movie stacks collected with 40-degree tilt. Movie stacks were imported and processed using the cisTEM processing pipeline (Grant et al., 2018). Briefly, motion-corrected and aligned micrographs with CTF fits better than 4.5 Å were filtered prior to auto-picking with an inter-particle exclusion radius of 90 Å and particle radius of 60 Å, with a threshold height of one standard deviation above noise in cisTEM. One round of 2D classification was performed with 150 classes for particles from zero-degree tilt data and 100 classes for particles from 40-degree tilt datasets. Classes were manually inspected and selected for ab initio model generation, followed by 3D auto-refinement. To improve the quality of the reconstruction, a mask excluding the peripheral IgM Cμ2 was applied, followed by 3D-classification with 5 classes. One class containing the best map features was selected for iterative 3D refinement. Additional focused refinements around IgM-Fc Cμ4 and SC-D1, D3-5 or IgM-Fc1 and SC-D1 domains were performed to improve map quality in those regions. Focus refined maps were combined using phenix.Combine_focused_maps (Liebschner et al., 2019). Local resolution was determined using ResMap (Kucukelbir et al., 2013) and rendered on a linear scale from 2.6-5.0 Å in UCSF Chimera (Pettersen et al., 2004).

### Model Building, Refinement, and Validation

Homology models of human IgM Cμ3 and Cμ4 domains were built based on the NMR structure of murine Cμ3 (PDB 4BA8) and crystal structure of murine Cμ4 (PDB 4JVW) using SWISS-MODEL (Waterhouse et al., 2018). Homology models of IgM tailpiece segments were similarly generated from the cryoEM structure of pentameric sIgA (PDB 6UEA). Homology models, along with the JC and individual domains of the SC from pentameric sIgA, were docked and rigid-body fit into the map using phenix.realspacerefine (Afonine et al., 2018). The model was subsequently inspected and further improved through iterative model building in Coot (Emsley et al., 2010) and real-space refinement in Phenix. Models were validated using phenix.validation_cryoem (Afonine et al., 2018) as well as EMRinger (Barad et al., 2015). Figures were generated using PyMOL (The PyMOL Molecular Graphics System, Version 2.2.2 Schrödinger, LLC) and UCSF Chimera (Pettersen et al., 2004) and ChimeraX (Goddard et al., 2017).

### Sequence Alignments and Conservation Analysis

Sequences for human IgM-Fc (342-576), IgA1-Fc (241-472), and IgA2-Fc (241-472) were aligned using Clustal Omega with default settings (Madeira et al., 2019), and the results rendered by conservation index on structure of IgM-Fc/JC in PyMOL.

## Acknowledgements

We thank Christine Tam, Yvonne Franke, and the Biomolecular Resource group for construct generation and the Research Materials Group for protein expression. We thank members of the Structural Biology department and the cryo-EM group for helpful discussions, including Alexis Rohou for advice on data processing and Jian Payandeh for helpful discussions on manuscript preparation.

## Data and materials availability

The cryo-EM map and coordinate model have been deposited to the Worldwide Protein Data Bank with PDB accession code 7K0C and EMDB code EMD-22591.

---

**Figure 1 - figure supplement 1:**
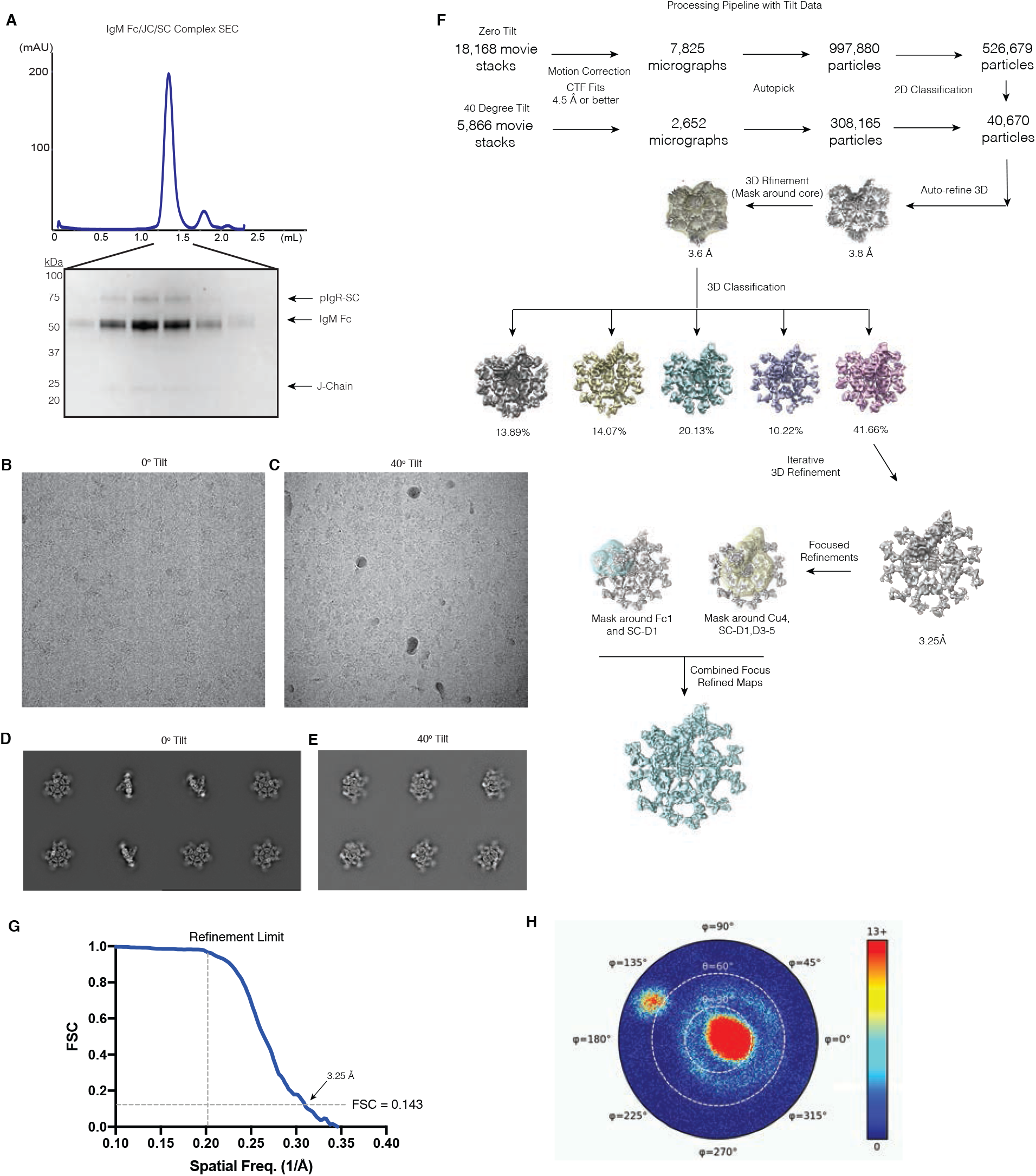
SIgM purification and cryo-EM data processing. (A) Representative size-exclusion chromatography trace (top) and reducing SDS-PAGE gel (bottom) of pentameric IgM-Fc/JC/SC complex. Representative motion-corrected and aligned cryo-EM images of samples imaged with zero degree (B) and 40 degree tilt (C) along with selected corresponding 2D class averages for zero degree (D) and 40 degree tilt (E) datasets. Summarized data processing pipeline along (F), along with FSC curve for the final reconstruction (G), with refinement limit indicated by the dashed vertical line and FSC=0.143 indicated by the dashed horizontal line. (H) Corresponding angular distribution plot for reconstruction shown in (G).

**Figure 1 - figure supplement 2:**
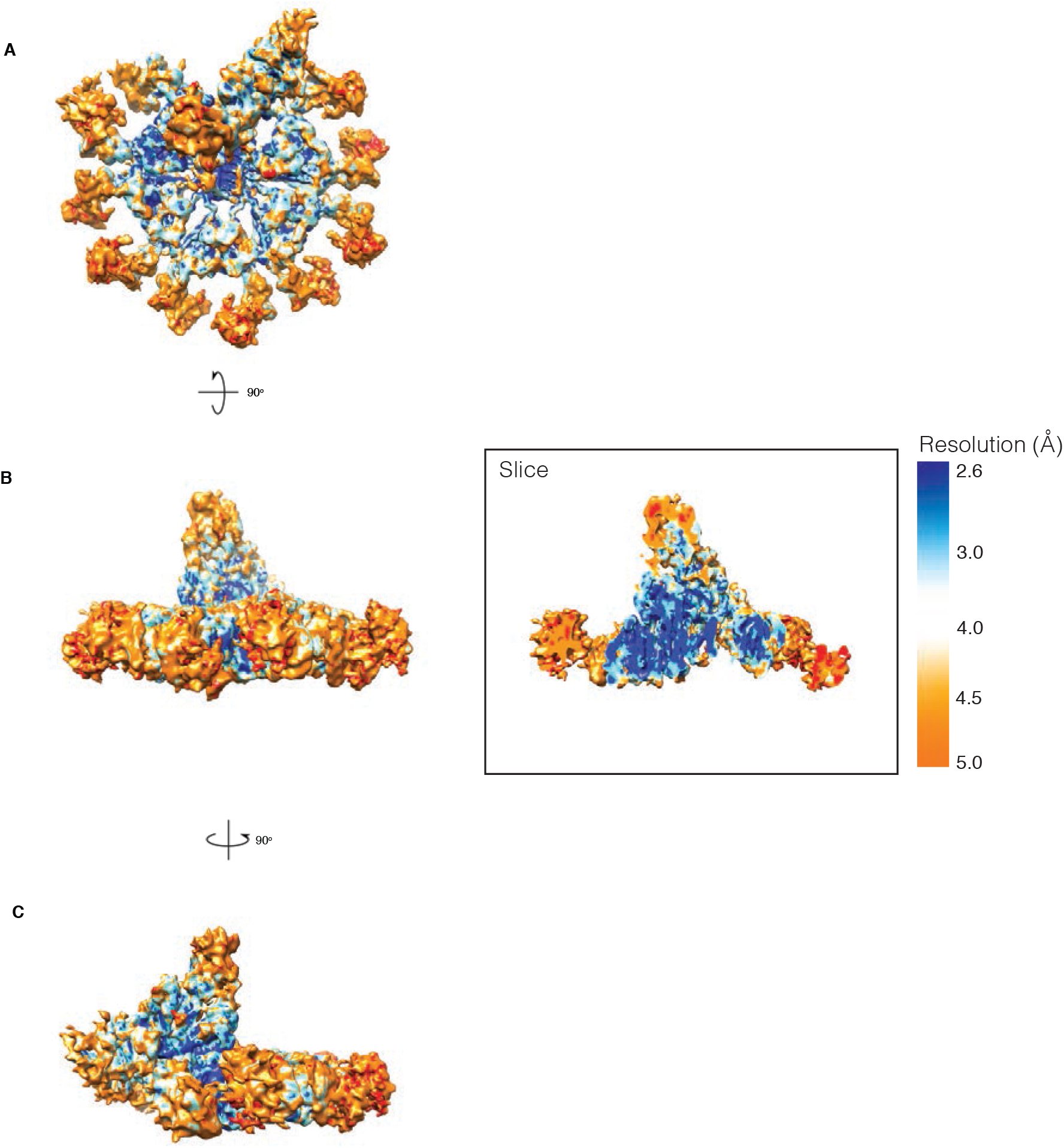
Local resolution of the sIgM reconstruction. Top (A), front (B), and side (C) views of the IgM reconstruction colored by local resolution from blue to red. Slice through SC-D1 and Fc interface from front view in (B) shown on right.

**Figure 1 - figure supplement 3:**
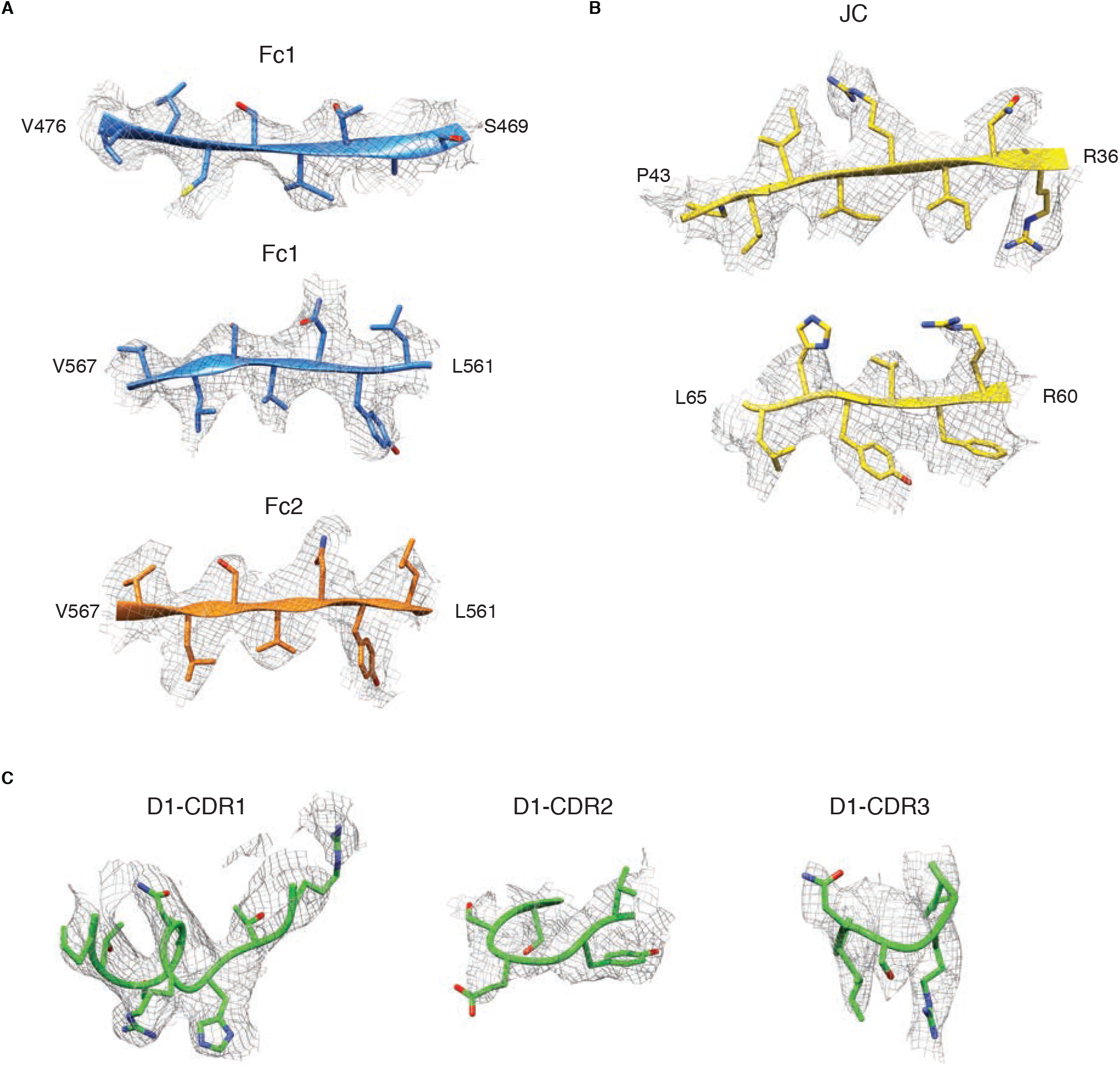
Cryo-EM maps of select regions of sIgM complex. (A) Density of β-strands corresponding to IgM Fc1 and Fc2. (B) Density corresponding to select regions of the JC. (C) Density corresponding to SC-D1 CDR1, 2, and 3.

**Figure 2 - figure supplement 1:**
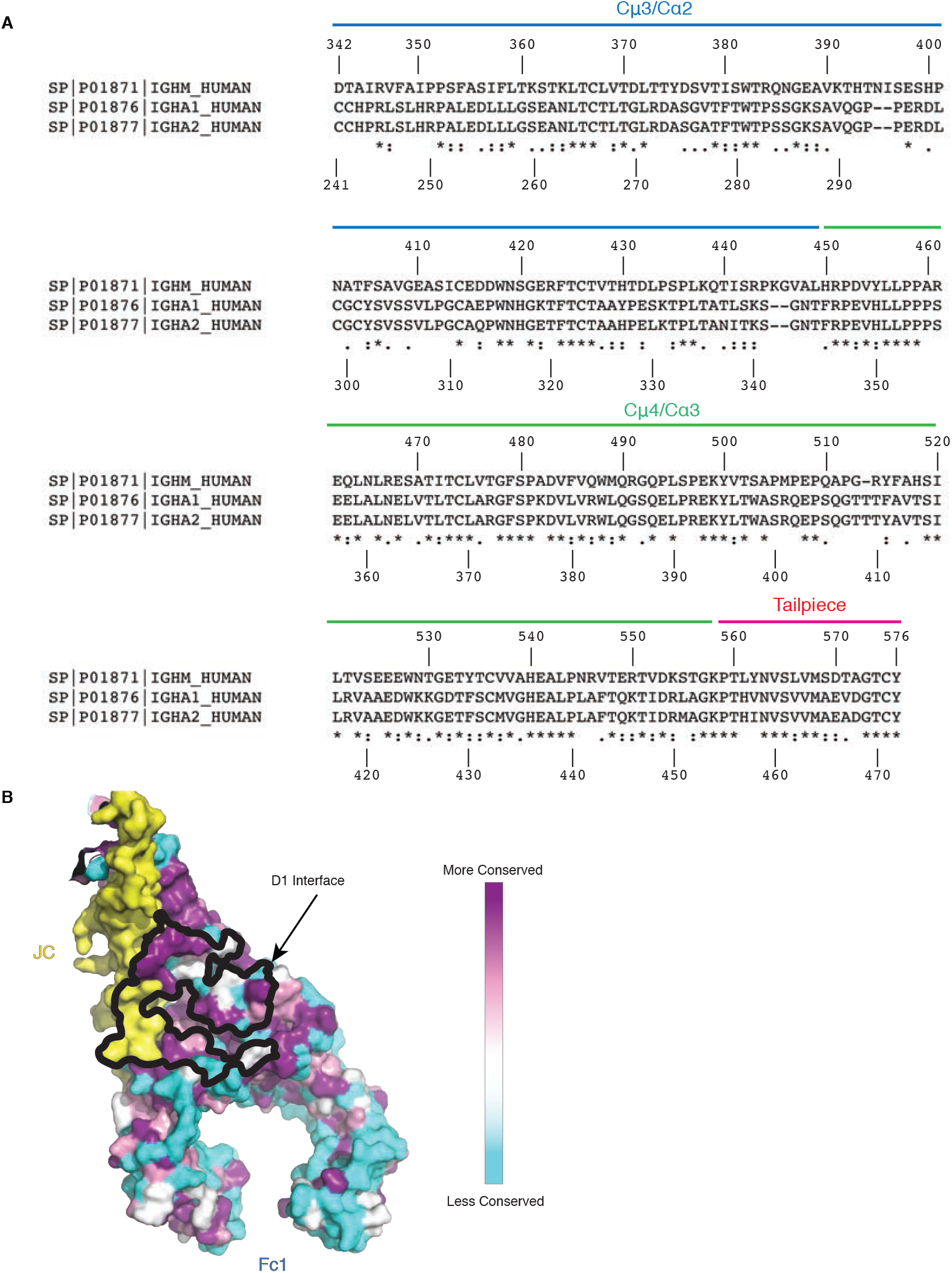
Sequence alignments and surface conservation of SC-D1 interface with IgM-Fc/JC. (A) Clustal Omega multiple sequence alignments of human IgM residues 342-576 (UniProt P01871), human IgA1 residues 241-472 (UniProt P01876), and human IgA2 residues 241-472 (UniProt P01877). (B) Top-down view of IgM-Fc/JC colored by sequence conservation with IgA-Fc with SC-D1 omitted for clarity. Interaction interface of SC-D1 is outlined on the surface in black. Conservation index: Identical residues (maroon); residues with highly similar chemical properties (pink); residues with weakly similar chemical properties (white); dissimilar residues (cyan).

